# Generative cell phenotyping with structured latent populations

**DOI:** 10.64898/2026.06.30.735507

**Authors:** Fanny Bodart, Adrien De Voeght, Frédéric Baron, Gilles Louppe

## Abstract

Flow cytometry produces high-dimensional single-cell protein measurements central to immunophenotyping and clinical monitoring. Yet analysis still relies largely on manual gating, which is labour-intensive, poorly reproducible, and ill-suited to large marker panels. Existing computational approaches address classification or discovery in isolation, treating cell-type identity as a post-hoc annotation rather than as part of the generative model itself. We present MARVIN, a semi-supervised variational autoencoder that encodes the assumption that cells organise into discrete populations with continuous intra-population variability through a Gaussian mixture prior in the latent space. Because each component represents a distinct cell population, classification, discovery, and density estimation emerge as complementary views of the same representation. On public benchmarks, MARVIN matches or exceeds existing methods using as few as 10% labelled cells. Trained exclusively on healthy samples, it identifies leukaemic cells through elevated reconstruction error, providing an unsupervised anomaly detection signal. On paired stimulation data, it maintains stable population assignments while capturing condition-specific shifts in abundance and marker expression at patient-level resolution. MARVIN is open-source and designed for local deployment, adapting to institution-specific panels and instruments.

## Introduction

Flow cytometry (FCM) or mass cytometry enables the profiling of individual cells through the expression of surface and intracellular markers. It is used routinely in the diagnosis and monitoring of haematological malignancies [1], the assessment of measurable residual disease (MRD) [2], immune exploration [3], and the discovery of new immune populations [4]. Modern panels now measure 20 to 40 markers simultaneously, yet analysis still relies on manual gating, the iterative two-dimensional selection of cell populations by expert operators. While disease-specific guidelines exist to standardize its use [5], the approach is labour-intensive, scales poorly to large panels, and introduces operator-dependent variability that limits reproducibility [6]. These limitations have motivated a sustained effort to automate cytometry analysis using computational methods [7, 8].

Existing computational methods differ in how they assign cell-type identity. Clustering methods, including Gaussian mixture models [9], sefl-organizing map as FlowSOM [10], and k-nearest-neighbor graphs as PhenoGraph [11], recover structure in the data, but the resulting clusters must still be mapped to populations by hand. Supervised classifiers [8, 12] annotate known populations accurately, but cannot discover populations absent from their training labels. Knowledge-driven generative models such as SCYAN [13] fix the populations in advance through an expert marker-to-population table. Integration-oriented variational autoencoders such as CytoVI [14] embed cells in a continuous latent space, but recover identity by clustering that embedding afterwards. In all of these, cell-type identity is established outside the model, fixed in advance or recovered after the fact, and never learned as part of the generative process.

A more natural approach is to build cell populations directly into the model. The biological prior is straightforward: the immune system is composed of distinct cell populations, each exhibiting continuous phenotypic variation around a characteristic marker profile. A model that mirrors this structure, first determining which population a cell belongs to and then accounting for variation within that population, can perform classification, population discovery, and anomaly detection within a single framework without requiring separate tools or post-hoc analysis steps. Here we introduce MARVIN (Mixture-based Variational Autoencoder for Representation and Variation in Immune Networks), a deep generative model in which each cell population is explicitly represented as a distinct component. The model learns both to generate realistic cells from each population and to infer, given observed marker expression, which population a cell belongs to and how it varies within it. A small number of expert-annotated cells (as few as 10% of the dataset) is sufficient to anchor the model’s populations to known cell types, while additional components can capture previously unannotated or rare subsets. When the model encounters cells that do not fit any learned population, it flags them through poor reconstruction, providing an automatic signal for detecting anomalies such as residual leukaemic cells.

We evaluate MARVIN on three public mass cytometry benchmarks and one in-house clinical dataset from the University Hospital of Liège, demonstrating competitive classification from minimal expert annotation, recovery of masked rare populations, detection of leukaemic cells as anomalies, and stable population tracking across paired experimental conditions at patient-level resolution.

## Results

### Model overview

MARVIN is a generative model: it describes how each measured cell is produced, as a hierarchical variational autoencoder. A cell is assumed to belong to one population (a cell type) and to vary continuously around that population’s typical marker profile. It is produced in three steps: its population *c* is drawn from a prior probability over the cell types (*p*_*π*_(*c*)); a phenotype within that population is set by drawing around the population’s characteristic profile (a Gaussian *p*_*β*_(**z** | *c*) in a latent space **z**); and that phenotype is turned into the measured marker intensities **x** (a decoder *p*_*θ*_(**x** | **z**)). The discrete population fixes what a cell is, the continuous code fixes how it varies within that population.

Reading a cell’s markers, MARVIN infers its population and its phenotype. It returns a probability for each population (*q*_*ω*_(*c* | **x**)): a peaked, confident answer is a clear cell type, a spread-out answer flags a cell as ambiguous or new and worth a manual look. A small set of expert-labelled cells ties the populations to known cell types (Fig. 1).

**Figure 1:**
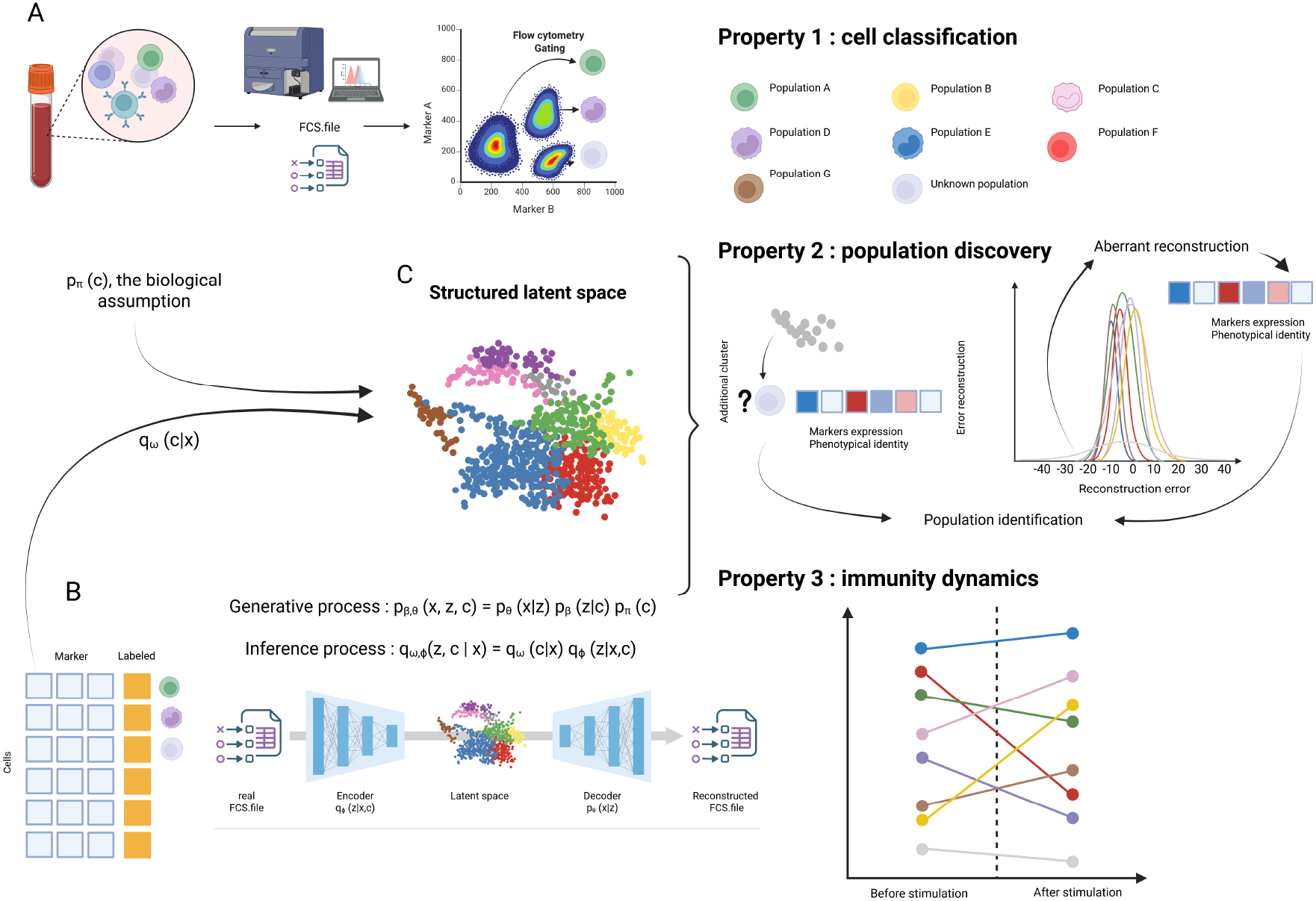
Overview of MARVIN. (A) Flow cytometry of a clinical sample yields an FCS file of per-cell marker intensities, conventionally analysed by manual gating. (B) MARVIN is a variational autoencoder whose latent space is a Gaussian mixture, one component per cell population. The mixture is shaped by two ingredients: a categorical prior *p*_*π*_ (*c*) over populations (their number and expected proportions) and a supervised encoder *q*_*ω*_ (*c*| **x**) that ties components to known cell types from a few labels. (C) The structured latent space supports three uses of a single model. *Classification*: each component carries a population’s phenotypic signature, so cells from a new sample on the same panel are annotated automatically. *Discovery*: spare components, or cells with high reconstruction error, capture populations absent from the labels. *Dynamics*: identity (*c*) is separated from state (**z**), so population assignments stay stable across conditions while their proportions are tracked.

### A small fraction of labels structures the latent space

Cytometry measurements carry technical variation from the instrument, the marker panel, and the acquisition conditions, which differ between centres. Left unchecked, a model can learn these site-specific patterns instead of biology. MARVIN guards against this by using a small fraction of expert-labelled cells to tie its populations to true cell types.

To determine how little labelling suffices, we trained on the AML mass-cytometry dataset [15] across labelled fractions from 0 to 100% (Fig. 2; Online Methods). The effect is clearest in the latent space, the model’s internal map of the cells. Even without labels, cells already form clusters, but phenotypically close populations overlap and are not resolved. A small fraction of labels separates them: the two NK subsets that differ only in CD16, for instance, are merged without supervision and cleanly resolved at 10%.

**Figure 2:**
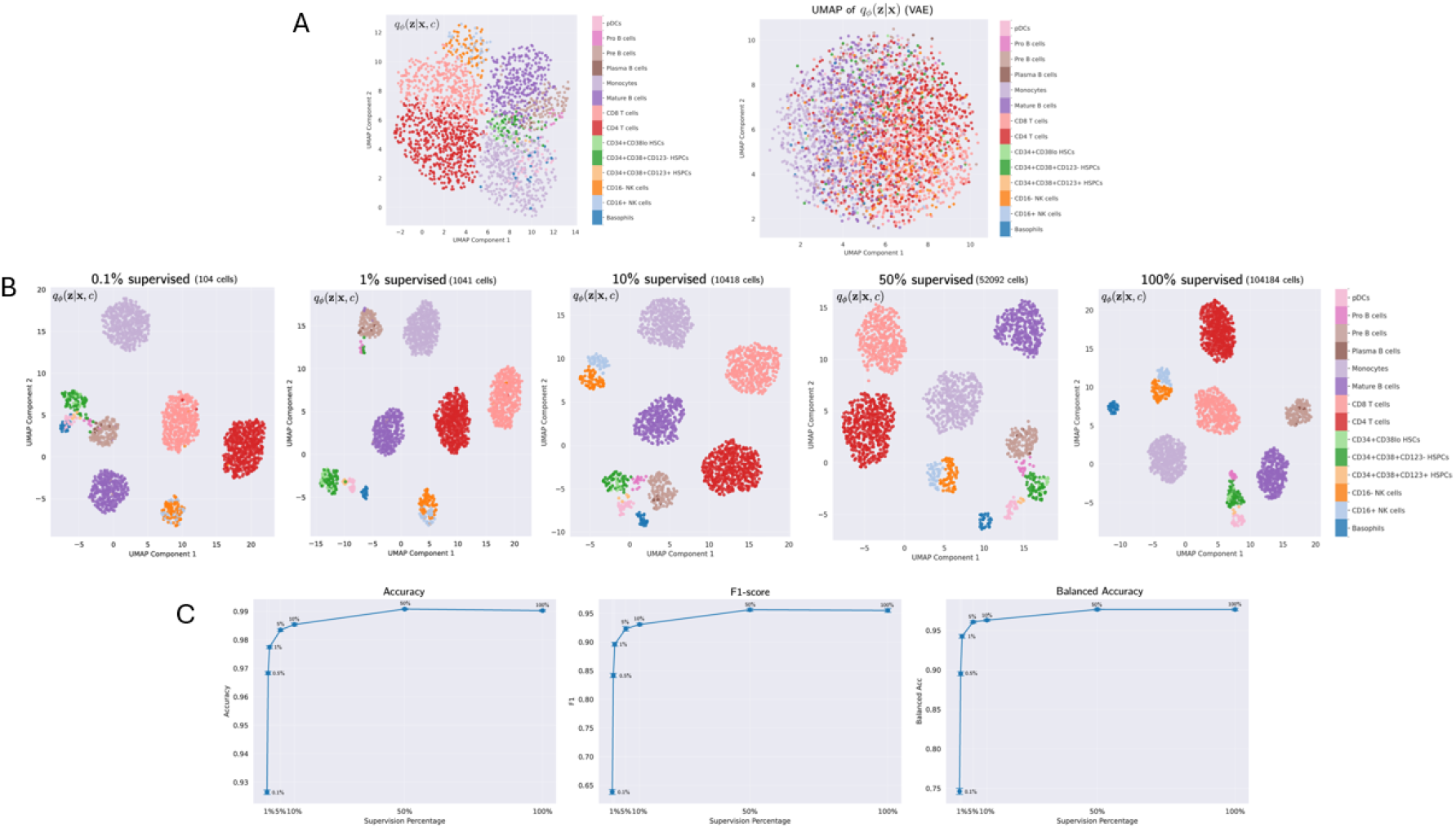
A small fraction of labels structures the latent space (AML). (A) UMAP [29] of the latent space without supervision, coloured by ground-truth label. Even unsupervised, MARVIN’s mixture structure (left) groups cells of the same type, unlike a standard VAE (right), whose latent space is unstructured. (B) The same 2,000 cells as supervision increases from 0.1% to 100%, coloured by ground-truth label. Phenotypically close populations that overlap at low supervision, such as CD16^+^ and CD16^*−*^ NK cells, separate cleanly by 10%. (C) Accuracy, F1, and balanced accuracy against supervision level: performance rises sharply with the first few labels and saturates near 10%.

We also tracked classification performance across these supervision levels. Accuracy is misleading here because the cell types are highly imbalanced, so it barely moves (0.92 to 0.99). Balanced accuracy and F1, which give equal weight to rare populations, rise from 0.75 to 0.95 and from 0.65 to 0.95. Ten percent of labels is therefore enough to recover the rare populations.

We, then, evaluate how supervision influenced accuracy on other datasets, especially bigger datasets. In the POISED dataset, millions of cells are present. When dataset is becoming bigger, the percentage of supervision decreases as there are enough absolute labelled cells to have a representation of the diversisty of populations. From 10% in AML, only 1% of labelled cells were enough to achieve high accuracy (Suppl. Fig. S1)

### Classification from few labels

Cells of the same type share a reproducible marker signature, so MARVIN should label them consistently. We tested this by using MARVIN to annotate cells and comparing its labels with expert manual gating. As a baseline we used SCYAN [13], a published deep generative model for cytometry annotation. The comparison spanned four datasets: three public mass-cytometry benchmarks (AML [15], BMMC [11], and the peanut-allergy cohort POISED [16]) and one in-house flow-cytometry dataset of bone-marrow samples from the University Hospital of Liège, under a shared preprocessing pipeline and 10% of cells labelled (Fig. 3; Online Methods).

**Figure 3:**
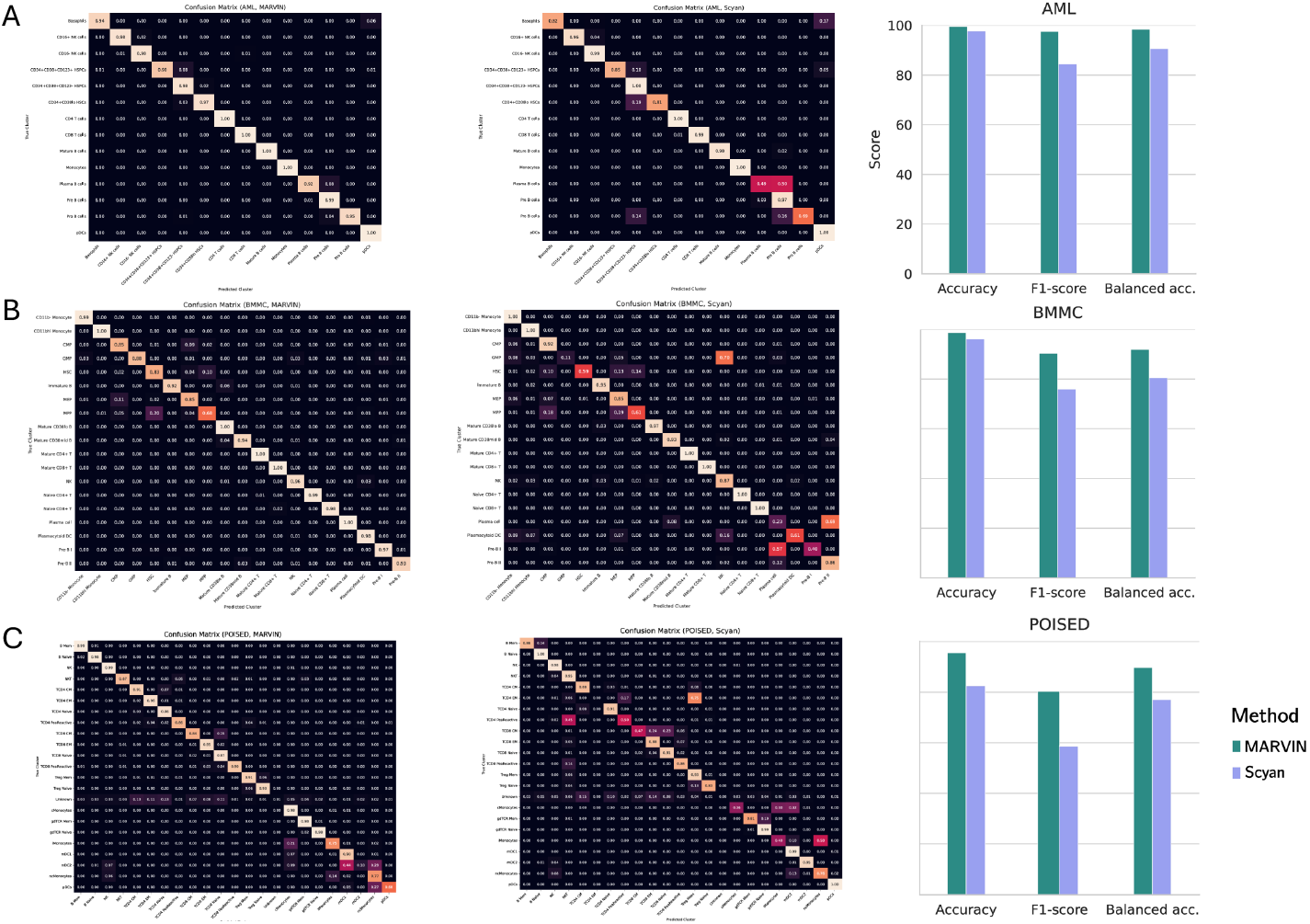
MARVIN matches or exceeds SCYAN on three public datasets. Confusion matrices against expert manual gating, with accuracy, F1, and balanced accuracy for MARVIN and SCYAN, on (A) AML, (B) BMMC, and (C) POISED. MARVIN’s matrices are more strongly diagonal, and its advantage over SCYAN is largest on the imbalance-aware metrics and on POISED, the most phenotypically complex dataset.

MARVIN matched or exceeded SCYAN on every dataset, most clearly on the imbalance-aware metrics. Balanced accuracy reached 0.98, 0.93, and 0.90 on AML, BMMC, and POISED (SCYAN: 0.89, 0.80, 0.78) and F1 reached 0.97, 0.92, and 0.80 (0.83, 0.75, 0.58), while accuracy, inflated by the class imbalance, reached 0.99, 0.99, and 0.95 (0.98, 0.96, 0.81). The margin was largest on POISED, the most phenotypically complex dataset, where MARVIN’s built-in population structure helps most. We also evaluate MARVIN under the same modality as it was performed with SCYAN that offer the possibility to pre-select markers of supposed importance. So reducing the panel to the markers most used in gating (14 of 32 for AML, 19 of 39 for POISED) left AML accuracy unchanged and lowered POISED accuracy slightly.

On the in-house dataset, MARVIN was trained on one labelled patient and tested on two others. It generalised well to the first (balanced accuracy 0.97, F1 0.95, accuracy 0.99) and less well to the second (0.92, 0.78, 0.76), consistent with inter-patient variation rather than model failure.

### Discovery of unannotated populations

Clinically important populations are often rare or undescribed: disease-associated subsets, new activation states, or cells missed by the gating strategy. MARVIN can be allowed more populations than are known; the spare components carry no label and are free to capture cell groups that the labelled types do not account for.

To test this, we removed the labels of the CD4 peanut-reactive T cells, a population at 0.1% of the POISED cohort, and allowed *K* = 24 populations, three more than the 21 that remained labelled (22 gated populations, one hidden). MARVIN placed 95.9% of the unlabelled peanut-reactive cells in a single spare component and kept them distinct from the known populations. That component also drew in a few conventional CD4^+^ cells, as expected from their phenotypic closeness to the peanut-reactive ones. The other two spare components captured CD4 central-memory and naive subsets marked by CD25, CLA, and specific TCR chains. The spare capacity was used rather than left empty, as it improved reconstruction, so it captured real structure rather than over-splitting the data.

We sampled 3,000 cells from the model’s prior and embedded them with t-SNE [17]: the recovered peanut-reactive component formed its own region near the related CD4^+^ populations, while the other two spare components sat closer to their parent populations (Fig. 4). Distance in the latent space tracks biological distinctiveness.

**Figure 4:**
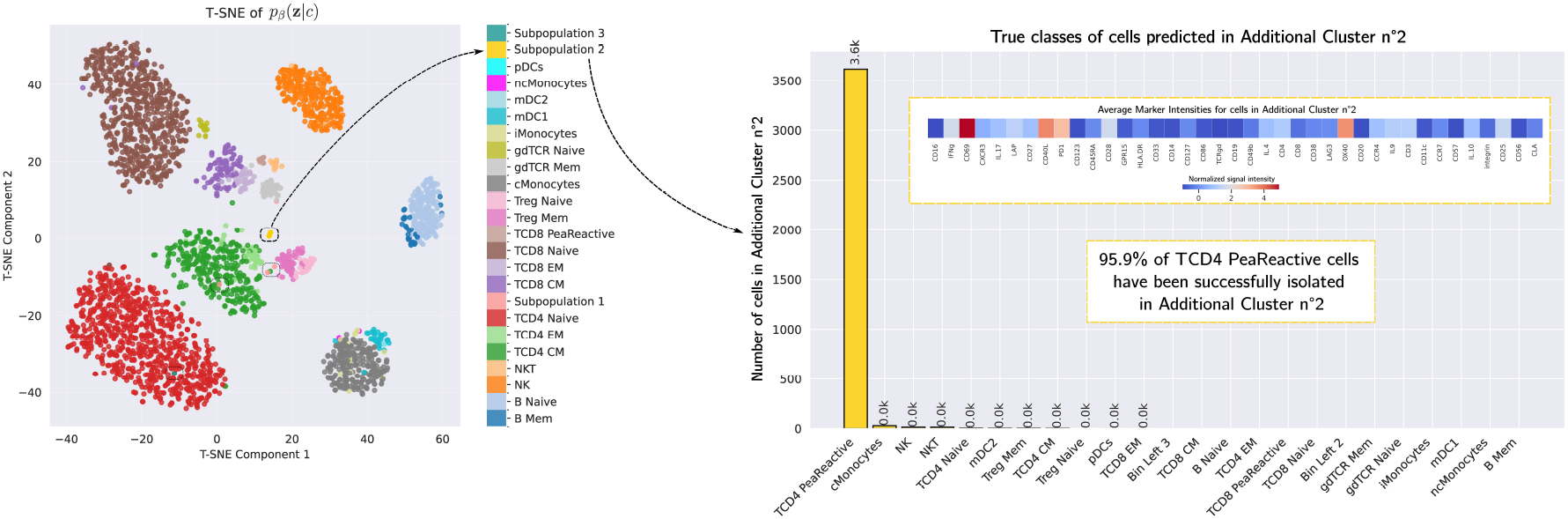
MARVIN recovers a masked rare population (POISED, *K* = 24). t-SNE of the latent space when three spare components are added and the CD4 peanut-reactive labels are withheld during training. One spare component (“new subpopulation 2”) collects 95.9% of the withheld CD4 peanut-reactive cells in a region of its own, near the related CD4^+^ populations, recovering a population the model was never shown.

### Detection of leukaemic cells as anomalies

Minimal residual disease monitoring asks whether a small fraction of leukaemic cells persists among many healthy ones [2]. Leukaemic cells can carry patient-specific marker combinations never seen in training, which makes rule-based gating unreliable. MARVIN instead scores each cell by how well it can reproduce it from the learned populations; a cell unlike anything it has seen reconstructs poorly and receives a high score (a high negative log-likelihood, NLL).

We trained MARVIN on healthy cells from the in-house dataset and introduced leukaemic cells only at test time. Healthy cells reconstructed well (NLL −6 to 0), while the leukaemic cells, absent from training, reached an NLL near 20, even though they made up under 0.2% of the test set (Fig. 5A). MARVIN therefore flags pathological cells without having been trained on any disease state. As a control, we then included leukaemic cells in training: their NLL narrowed to the healthy range, confirming that the high score reflects their absence from training rather than noise or class imbalance (Fig. 5B).

**Figure 5:**
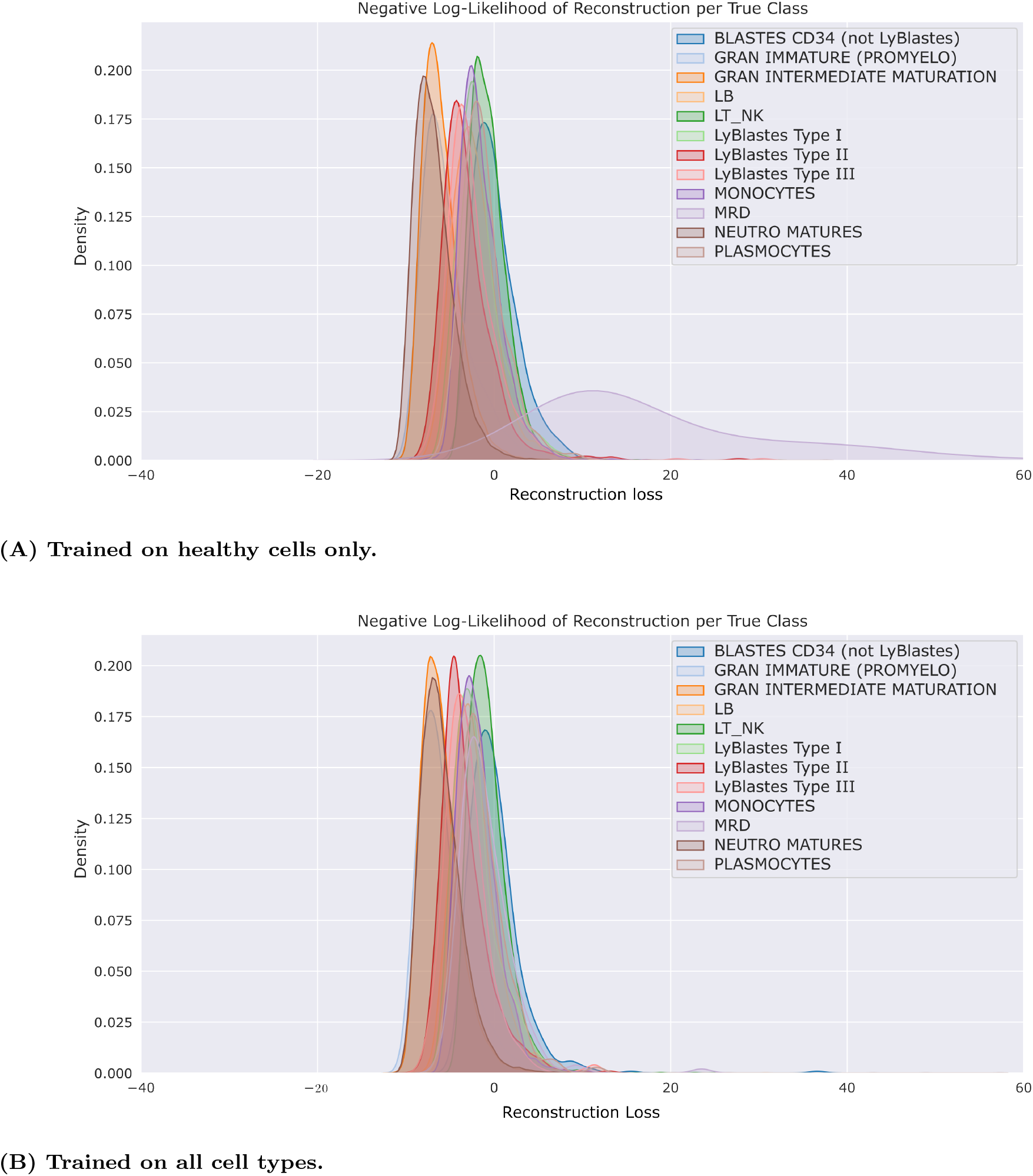
Reconstruction error separates leukaemic from healthy cells. Per-cell negative log-likelihood (NLL) under the decoder, by true class, estimated by kernel density on 500,000 cells. (A) Trained on healthy cells only, the model reconstructs them at low NLL but assigns markedly higher NLL to leukaemic (MRD) cells, flagging them as anomalies although it never saw them in training. (B) Trained on all cell types, the leukaemic NLL collapses into the healthy range, confirming that the high NLL in (A) reflects their absence from training rather than an inability to represent them.

### Tracking population dynamics across conditions

A clinically useful tool must also track how population composition changes across conditions or over time, for instance as the immune system responds to an antigen. We used the POISED dataset, which pairs a baseline and a peanut-stimulated sample from each of 15 peanut-allergic patients.

Across the cohort, stimulation shifted population proportions, and the CD4^+^ peanut-reactive population increased after exposure, as expected for an antigen-specific T-cell response (Fig. 6A). MARVIN assigned stimulated and unstimulated peanut-reactive cells to the same population despite their different activation markers (CD69, CD40L, OX40). The population assignment (*c*) stays fixed while the continuous code (**z**) absorbs the activation state, so MARVIN separates what a cell is from the state it is in (Fig. 6B).

**Figure 6:**
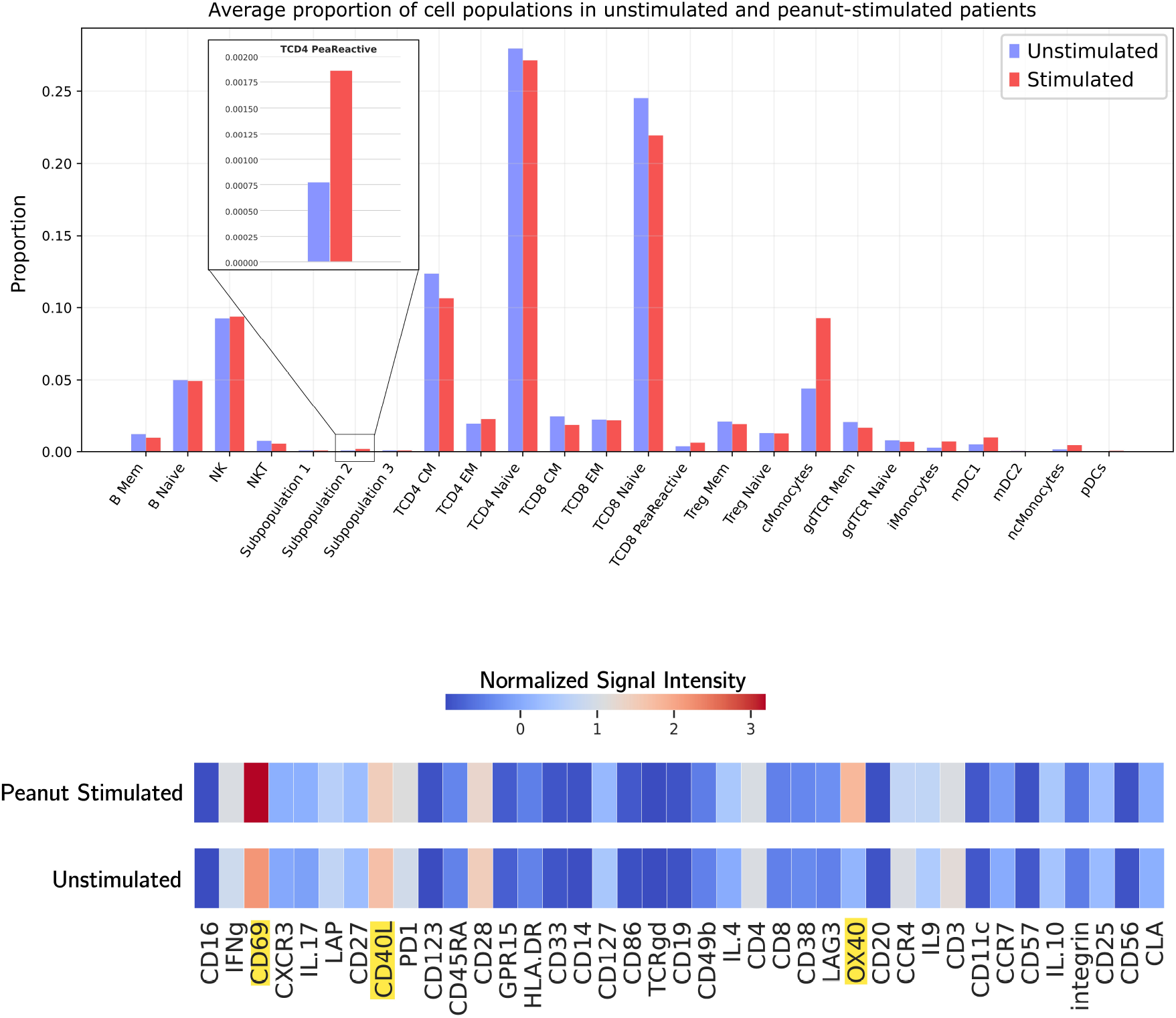
MARVIN tracks population dynamics while keeping identity stable (POISED). (A) Mean population proportions under unstimulated versus peanut-stimulated conditions, averaged across patients; the discovered CD4 peanut-reactive population (subpopulation 2, inset) increases after stimulation, as expected for an antigen-specific response. (B) Mean marker expression of CD4 peanut-reactive cells, stimulated versus unstimulated: CD69, CD40L, and OX40 rise with activation, yet MARVIN assigns both conditions to the same population, separating activation state from cell identity.

At the patient level, proportions varied widely at baseline and after stimulation. The CD4^+^ peanut-reactive population rose after stimulation in most patients, with a few limited or atypical responses, and classical monocytes rose in most patients as well (Fig. 7). MARVIN therefore supports both cohort-level and patient-level analysis while keeping population identity stable across conditions.

**Figure 7:**
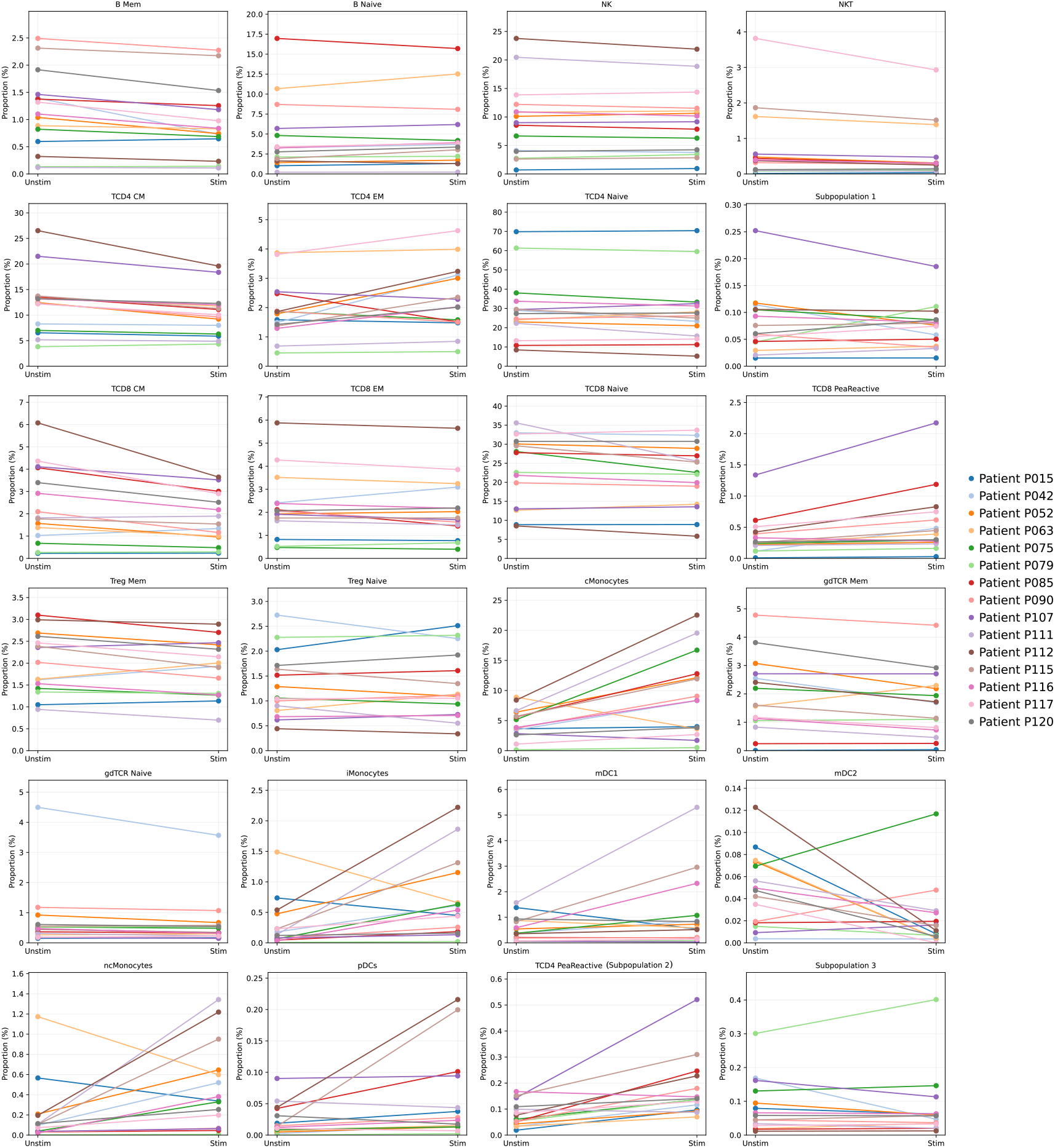
Patient-level population dynamics. For each population, the change in its proportion from the unstimulated to the peanut-stimulated condition, one line per patient. Responses are heterogeneous: the CD4 peanut-reactive population and classical monocytes rise in most patients, with a few limited or atypical responses.

## Discussion

We introduced MARVIN, a semi-supervised generative model for high-dimensional flow and mass cytometry that builds cell-type identity into the model itself. By structuring the latent space as a Gaussian mixture within a variational autoencoder, with one component per population, MARVIN serves classification, discovery, and density estimation from a single inference procedure. Recovery of rare or unannotated subsets and assessment of distributional shifts across conditions follow from the same representation.

### Relation to existing methods

Methods for automating cytometry differ in how much biological knowledge they build in, and how. Unsupervised clustering, such as FlowSOM [10], groups cells by marker expression without prior knowledge: a self-organising map followed by hierarchical clustering partitions the data into coherent groups. The clusters are cheap to compute and scale to millions of cells, but they must be interpreted and matched to known cell types after the fact, and nothing associates a cluster with a biological identity during training. Rare populations, those under 1% of the data, suffer most, because the model has no signal that they are worth preserving.

SCYAN [13] replaces manual labels with a knowledge table that states which markers each population expresses, and maps marker expression through a normalising flow into a latent space constrained by that table. It needs no cell-by-cell labels, but the table itself encodes expert decisions about which populations exist and how they are defined, so the supervision moves from cell-level annotation to population-level prior knowledge rather than disappearing. A marker left unspecified for a population is sampled from a uniform prior (*e*_*m*_ ~ *U* ([−1,1])) and treated as uninformative; in large panels many entries are undefined, so much of the available signal is discarded and the latent space is reliable only for the markers the table constrains. Markers outside the predefined strategy, even diagnostically useful ones, cannot contribute, and classification accuracy depends on how many markers the table specifies. Discovery is likewise a post-hoc, user-driven step: a population is selected, subdivided by external clustering of the latent space, and characterised there, rather than emerging from the generative objective.

CytoVI [14] is a generative VAE built for cross-dataset integration. It embeds cells in a continuous latent space, under a Gaussian prior or, when cell-type labels are supplied, a mixture prior whose modes are tied to the labels, and uses the generative model to impute missing markers, correct batch effects, and test differential abundance. The embedding carries a per-cell uncertainty, but cell-type identity is not encoded in it: cell types are assigned after training by clustering the embedding, so there is no per-cell confidence in the assignment itself. CytoVI is strong for harmonising studies, technologies, and panels, but for clinical uses such as MRD monitoring or rare-subset detection, which need per-cell assignment scores, this post-hoc annotation is a real limitation.

MARVIN differs from these methods by learning population identity during training rather than fixing or recovering it outside the model. Each mixture component is a cell population whose phenotypic signature is learned from labelled examples, not specified by a knowledge table or recovered by post-hoc clustering. MARVIN therefore adapts to local panels and instruments without a population-level knowledge table, while still using the structure that expert labels provide. Because the supervised signal guides the latent space during training rather than constraining it beforehand, every marker in the panel contributes to the representation, including those a knowledge table would leave unlinked. The discrete variable *c* gives per-cell uncertainty through the entropy of *q*_*ω*_(*c* | **x**), and the reconstruction objective supports anomaly detection and discovery, without the separate procedures these tasks require in SCYAN and CytoVI.

### Experimental findings

The experiments bear this out. With 10% supervision MARVIN annotated cells robustly, and performance saturated at modest labelling, so explicit latent structure lowers the need for manual annotation while keeping biologically coherent organisation. Supervision helped rare populations most, consistent with the cross-entropy term stabilising minority-class assignments. Discovery worked in two ways. Spare latent components recovered a masked CD4^+^ peanut-reactive population into a single emergent cluster, a partition that survived the supervisory constraints rather than fragmenting arbitrarily. Reconstruction likelihood gave an anomaly signal: trained only on healthy cells, MARVIN assigned high negative log-likelihoods to leukaemic cells (under 0.2% of the data), which fell back to the healthy range once those cells were included in training. MARVIN can thus flag phenotypes absent from the training distribution, supporting anomaly detection and MRD monitoring that neither SCYAN nor CytoVI address in one framework.

Beyond static classification, MARVIN held population identities stable under perturbation while tracking shifts in abundance and activation markers. In POISED, peanut-reactive CD4^+^ T cells stayed in one latent component at baseline and after stimulation despite changing activation markers, so the learned signature is not tied to a single activation state, which enables longitudinal and paired-condition analyses at cohort and patient resolution.

### Limitations

The framework has limits. Each component is Gaussian, which may not capture non-elliptical or multimodal phenotypes in very heterogeneous populations. The number of components must be set in advance, and principled model selection needs further work. Configuration to local panels is supported, but transfer across substantially different marker sets remains open, and is partly what CytoVI targets. Anomaly detection through reconstruction depends on how representative the training data are, and should be read alongside biological validation.

### Deployment and outlook

For deployment, MARVIN needs only a small well-labelled reference cohort and then applies to further samples acquired under the same panel and settings, which suits inter-centre standardisation: train on a reference dataset centrally, deploy locally across sites. It runs on conventional fluorescence and spectral flow cytometry as well as mass cytometry. Each inferred population also has an interpretable signature: the average marker expression of its assigned cells, in the panel’s own markers. This lets a discovered population be matched to known gating definitions and, in principle, isolated for downstream functional assays.

Structured mixture-based variational models offer a flexible basis for cytometry analysis, joining annotation, discovery, and distributional modelling in one probabilistic framework. With limited supervision, MARVIN shows how biologically informed latent structure improves robustness, interpretability, and sensitivity to rare or emerging populations in high-dimensional single-cell protein data.

## Online Methods

### The MARVIN model

MARVIN is a hierarchical variational autoencoder [18] for cytometry data. The latent space is a Gaussian mixture: the immune system is assumed to comprise *K* cell populations, a discrete variable *c* selects the population, and a continuous code **z** captures variation within it. This construction follows the semi-supervised M2 model [19] and Gaussian-mixture and clustering variational autoencoders [20, 21].

### Problem definition

We observe *N* cells, each a vector of *M* marker intensities **x**_*i*_ ∈ ℝ^M^. A subset of cells carry an expert label *c*_*i*_ ∈ {1, …, *K*} giving their population; the rest are unlabelled. We want a model that assigns every cell to one of *K* populations, represents the continuous variation within each population, and flags cells that match no population. We cast this as learning a generative model *p*(**x, z**, *c*) together with an approximate posterior *q*(**z**, *c* | **x**), fitted by maximising a variational lower bound on the data likelihood with an auxiliary supervised term on the labelled cells.

### Generative process

The joint distribution of marker intensities **x**, latent code **z**, and population *c* factorises as

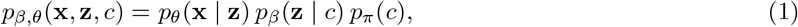

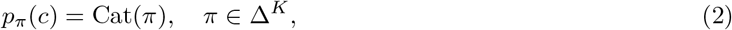

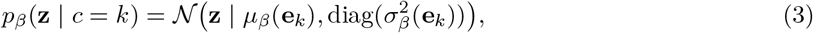

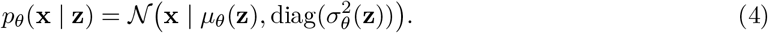

Here **e**_*k*_ is the one-hot encoding of population *k*, and 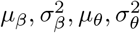 are computed by neural networks. Given **z**, the observation **x** is independent of *c*, so *c* serves only to structure the latent space.

### Inference process

We approximate the true posterior *p*(**z**, *c* | **x**) by a variational distribution that factorises as

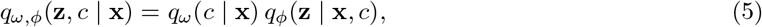

with a categorical posterior over populations,

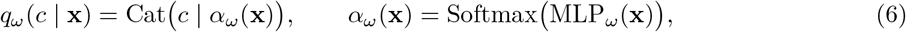

and a Gaussian posterior over the latent code,

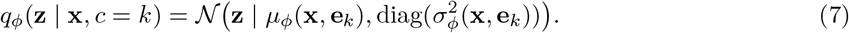

The categorical posterior *q*_*ω*_(*c* | **x**) is the classifier head: it predicts the population directly from **x**, and the softmax, rather than an argmax, preserves uncertainty for ambiguous cells and allows soft assignment across populations.

### Learning process

For an unlabelled cell, the evidence lower bound on log *p*(**x**) is

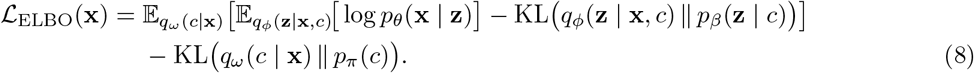

The three terms reward faithful reconstruction, keep each population’s latent code close to its Gaussian prior, and keep the inferred population distribution close to the categorical prior. For a labelled cell with label *c*^⋆^, a supervised cross-entropy term −log *q*_*ω*_(*c*^⋆^ |**x**) trains the classifier head directly. Training minimises this cross-entropy on the labelled cells X_*L*_ minus the ELBO over all cells X, with no relative weighting,

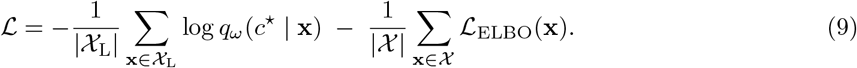

The mixture prior drives separation between populations, while the conditional encoder *q*_*ϕ*_ captures variation within each.

### Implementation

MARVIN is implemented in PyTorch [22]. Every head (*q*_*ω*_, *q*_*ϕ*_, *p*_*β*_, *p*_*θ*_) is a residual multi-layer perceptron of two residual blocks [23], with hidden layers eight times wider than the input and dropout [24] (rate 0.3) at the end of each block; self-attention and graph modules are not used, given the high-dimensional, non-spatial nature of cytometry data. The latent dimension *D* is set per dataset over {16, 32, 64}, with *D* = 64 for the more complex datasets.

We train with AdamW [25, 26], fixing *β*_*1*_ = 0.9 and setting *β*_*2*_ = 0.98 at the large batch sizes used for million-cell datasets, where the default 0.999 made the loss spike. The learning rate warms up linearly from 10^−5^ to 2 × 10^−3^ and is then halved every 10 epochs, over 30 epochs of training; the batch size is 64 for datasets under 500,000 cells and 1024 for larger ones. We clamp 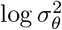 to [−6, 3], which keeps 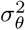 in roughly [0.002, 20] to avoid instabilities and overconfidence.

### Datasets

We used three public datasets and one in-house dataset.

The AML dataset [15] comprises 104,184 manually gated cells from bone marrow, with a panel of 32 markers and 14 populations. The BMMC dataset [11] comprises 61,725 bone-marrow mononuclear cells, with 13 markers and 19 populations. The POISED dataset [16] comprises 4,178,320 peripheral-blood mononuclear cells, with 39 markers and 22 gated populations; it contains 30 samples from 15 patients, one before and one after peanut exposure, and 982,741 cells were left ungated. The populations are highly unbalanced (Table 1): several, including the CD4 and CD8 peanut-reactive T cells of interest, fall below 1% of the data. We removed the ungated events for most experiments as no control could be possible.

**Table 1.** Cell populations from the POISED dataset.

| **Population** | **Proportion (%)** |
| --- | --- |
| Unknown (Ungated) | 23.52 |
| TCD4 Naive | 21.13 |
| TCD8 Naive | 17.81 |
| TCD4 CM | 9.12 |
| NK | 7.21 |
| cMonocytes | 5.78 |
| B Naive | 3.78 |
| TCD8 CM | 2.01 |
| gdTCR Mem | 1.40 |
| Treg Mem | 1.38 |
| TCD8 EM | 1.31 |
| TCD4 EM | 1.23 |
| Treg Naive | 1.13 |
| B Mem | 0.84 |
| mDC1 | 0.55 |
| gdTCR Naive | 0.48 |
| NKT | 0.44 |
| iMonocytes | 0.32 |
| TCD8 PeaReactive | 0.25 |
| ncMonocytes | 0.18 |
| TCD4 PeaReactive | 0.09 |
| mDC2 | 0.03 |
| pDCs | 0.03 |

The in-house MRD dataset comprises anonymised bone-marrow samples from three healthy donors, covering the major normal populations, together with leukaemic cells from four patients. Of 6,667,147 cells, 6,656,925 are healthy and 10,222 are leukaemic, so the leukaemic cells are a small minority. The data span 12 cell populations (11 healthy and one leukaemic) and 8 markers, and were preprocessed as for the public datasets (including an asinh transform [11, 13, 15, 27]).

### Experimental setup

#### Classification

All classification experiments used 1 to 50% of cells as labels, and we report accuracy, balanced accuracy, and *F*_*1*_-score against expert manual gating. As a baseline we ran SCYAN [13], a normalising-flow model that encodes a knowledge table of expected marker expression per population, under the same preprocessing pipeline. We implemented all the procedures as extensively describe on the repository. Briefly, we load the knowledge table (containing the phenotype of each cell population). 1 for positive expression of a marker, −1 if negative expression, NA if data is missing. Expression cal also be 0 or 0.5 for mid or low expression, respectively.). We used the standard parameters proposed in the SCYAN source code. We applied three metrics. Accurary is the ratio between correct prediction and total predictions. It is a global metric. Balanced accuracy compute the average recall for each class. It is more appropriate for imbalanced dataset. Formally balanced accurary is

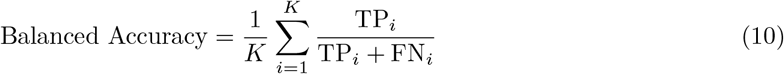

*F*_*1*_-score is the harmonic mean of precision and recall [28]. For imbalanced data, the true positive rate (recall) for the minority of class is an important goal. However, this improvement can increase the number of false positives and so, lowering precision.

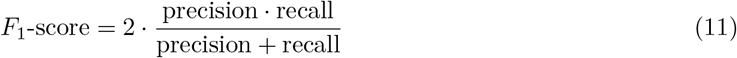

Where precision is

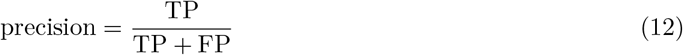

and recall is

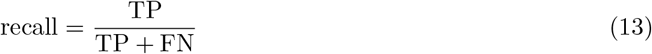

#### Discovery

Discovering unknown or shifted populations matters clinically: pathological cells can change phenotype during clonal evolution and escape a fixed manual-gating strategy. To test whether MARVIN recovers a population it was never shown, we used the POISED dataset (22 gated populations) and masked the labels of the CD4 peanut-reactive T cells, a population at about 0.1% of the data that is phenotypically close to other CD4 T cells. With these labels removed, the classifier head cannot help and the population can be recovered only through the unsupervised reconstruction. We fixed the prior *p*_*π*_(*c*) before training, giving one spare component a low weight (0.1%) to encourage it to capture a rare population, while the remaining components matched the empirical proportions of the labelled populations. We set *K* = 24, three more than the 21 populations that remained labelled, leaving three spare components for unannotated structure.

#### Anomaly detection

To test reconstruction-based anomaly detection, we trained MARVIN on the healthy cells of the in-house dataset, fixing *K* = 11 (the number of healthy populations), and introduced leukaemic cells only at test time. As a baseline, a second model was trained on the full dataset, healthy and leukaemic, with all labels kept. We scored each cell by the negative log-likelihood (NLL) of the decoder *p*_*θ*_(**x** | **z**), a feature-wise Gaussian: well-represented populations give a narrow, low NLL, while poorly represented or unseen ones give a broad, high NLL.

#### Population dynamics

We applied a trained MARVIN to the paired POISED samples and compared population proportions between the baseline and peanut-stimulated conditions, at both cohort and patient level. For the recovered peanut-reactive population, we also compared mean marker expression between the two conditions.

## Code and data availability

Code is available at https://github.com/fannybdt/MARVIN.git. The public dataset AML is available at https://figshare.com/articles/dataset/levine-CyTOF-annotated-celltypes-fcs/19867114. The original URL for the public datasets BMMC and poised are no longer available and datasets were donwload from the repository of SCYAN https://github.com/prism-oncology/scyan/blob/main/scyan/data/datasets.py. In-house data are available from the corresponding author on reasonable request.

## Algorithm 1

Training and generation with MARVIN

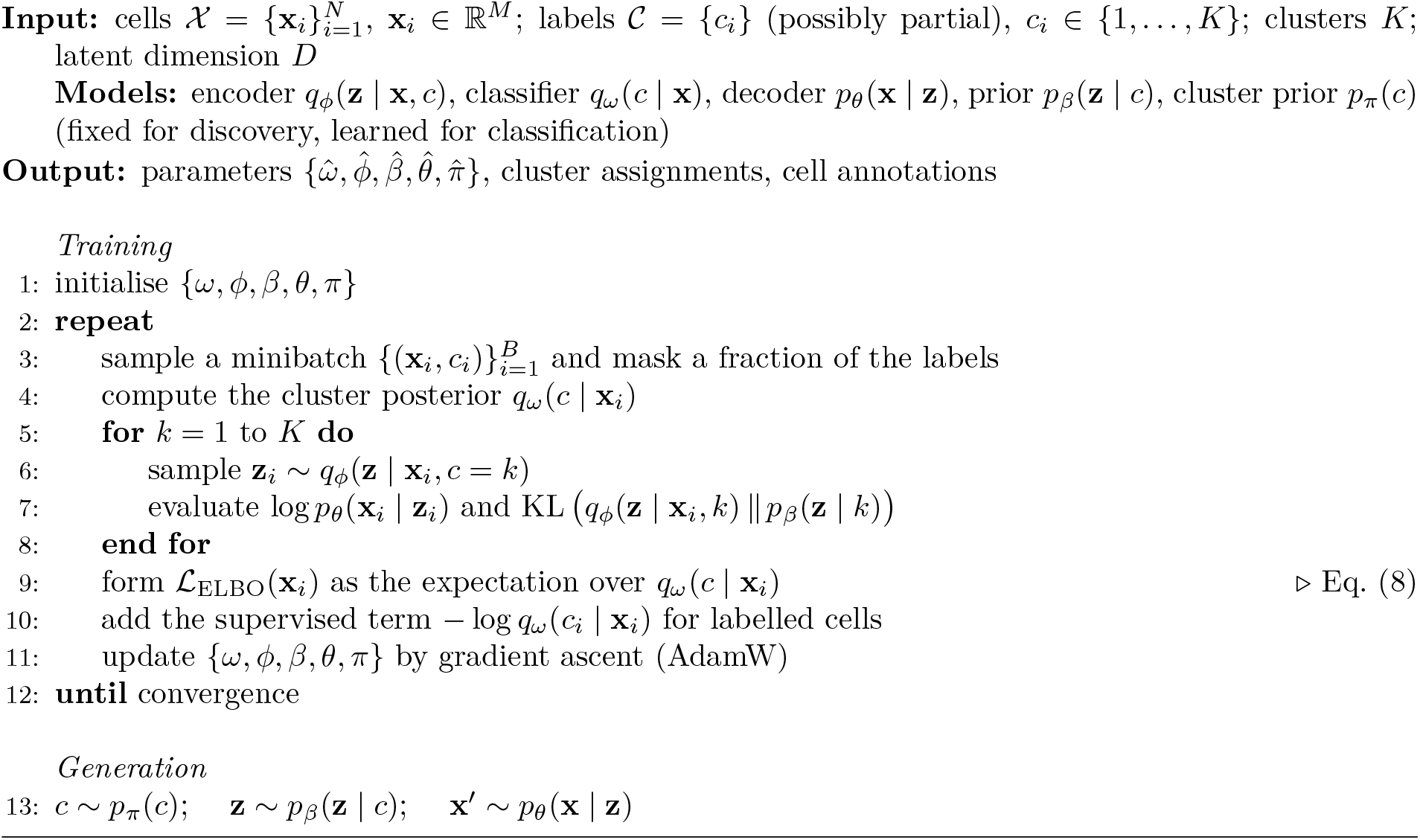

**Figure S1:**
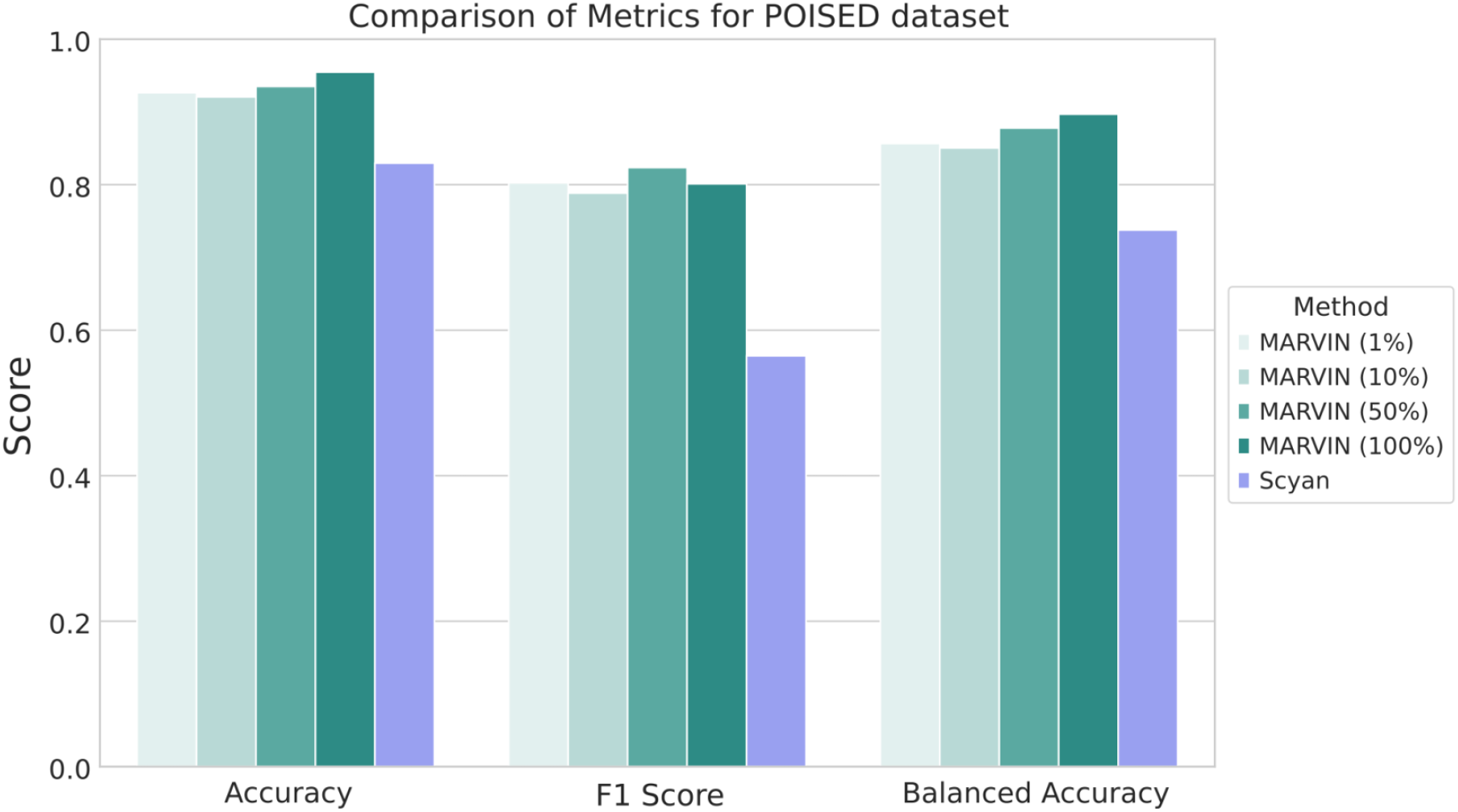
Comparison of Metrics for POISED dataset. The importance of supervision in a dataset of millions of cells remains important, however, even a small supervision of 1% is enough to achieve major performance across all the scores.

